# Genetic knockdown and knockout approaches in Hydra

**DOI:** 10.1101/230300

**Authors:** Mark Lommel, Anja Tursch, Laura Rustarazo-Calvo, Benjamin Trageser, Thomas W. Holstein

**Author notes:** contributed equally (Mark Lommel, Anja Tursch). Correspondence and requests for materials should be addressed to T.W.H.

## Abstract

Hydra is a member of the Cnidaria, an ancient phylum at the base of metazoan evolution and sister group to all bilaterian animals. The regeneration capacity of Hydra, mediated by its stem cell systems is unparalleled in the animal kingdom. The recent sequencing of the Hydra genome and that of other cnidarians has drawn new attention to this well-known model organism. In spite of this, the establishment of methods to manipulate gene expression in Hydra have remained a major challenge. Here we report a CRISPR-Cas9 based targeted mutation approach as well as an optimized, reproducible strategy for the delivery of siRNAs. Both approaches are based on a refined electroporation protocol for adult Hydra polyps. We demonstrate that these strategies provide reliable genetic interference with target gene expression, facilitating functional studies and genome editing in Hydra.

## Introduction

The freshwater polyp Hydra was the first animal used in studies of experimental developmental biology (Bosch et al., 2010; Technau and Steele, 2011; Trembley, 1744) and is famous for its almost unlimited regeneration capacity and longevity (Schaible et al., 2015). Hydra is member of the ancient phylum Cnidaria that exhibits a simple gastrula-like body plan with primary radial symmetry and two germ layers (Steele, 2012; Steele et al., 2011). Recent molecular phylogenies have revealed that cnidarians are the sister group to all bilaterian animals (Simion et al., 2017; Technau and Steele, 2011). This makes them an important model organism for understanding the origin of bilaterian body axes, the centralized nervous system, and the third germ layer (Holstein, 2012; Holstein et al., 2011; Steele et al., 2011; Technau and Steele, 2011). Hydra possesses three stem cell lines: the ectodermal stem cells give rise to ectodermal epithelial cells, endodermal stem cells to endodermal epithelial cells, and interstitial cells differentiate into neurons, nematocytes, germ cells, and gland cells (David, 2012; Steele, 2002; Steele et al., 2011).

In order to study the dynamics and function of genes in Hydra pattern formation and stem cell homeostasis several tools have been developed during the last years. Beside pharmacological treatment in analogy to vertebrate systems, the establishment of new molecular techniques facilitated the analysis of gene regulatory networks (GRN) and signaling pathways in the context of cell differentiation and patterning, making Hydra an attractive model for many experimental biologists.

A long-awaited breakthrough has been the introduction of foreign plasmid DNA into Hydra embryos by injection, eventually yielding stably expressing transgenic animals (Wittlieb et al., 2006). This technique made it possible to study the function of particular genes including their cis- and trans-regulatory elements (Nakamura et al., 2011). The introduction of plasmid DNA encoding for expression cassettes can also be applied for gain of function studies in adult polyps. For instance, electroporation-mediated delivery of the Nodal-related (Ndr) gene under control of the Hydra actin promoter into adult Hydra tissue resulted in increased budding rates (Watanabe et al., 2014).

For loss of function studies, the RNA interference (RNAi) approach (Fire et al., 1998) turned out to be most promising. Whereas morpholino-mediated knockdown of endogenous transcripts was only feasible in marine cnidarians (Momose et al., 2008; Rentzsch et al., 2008) the silencing of genes using double-stranded RNA (dsRNA) revealed to be efficient also in Hydra as shown by Lohmann and coworkers (Lohmann et al., 1999). Here, the introduction of dsRNA was mediated by electroporation.

In an alternative approach, HyAlx dsRNA was locally electroporated into early buds (Smith et al., 2000) or by introducing agarose pieces soaked with dsRNA-expressing bacteria into the gastric cavity (Miljkovic-Licina et al., 2007). The agarose approach is based on the feeding method developed for *C. elegans* (Conte and Mello, 2003) and planarians (Newmark et al., 2003), but requires more than two weeks treatment and leads to starvation and cell death (Technau and Steele, 2011). After the introduction of small interfering RNA (siRNA), which is more efficient and specific than long dsRNA (Elbashir et al., 2001), the RNAi approach has been further developed for Hydra (Watanabe et al., 2014). Although a significant downregulation of target genes was achieved in the ectoderm and endoderm, the damage induced by electroporation led to mortality rates greater than 50% within the first 24 hours post-electroporation. This hampered the development of RNAi to become a robust method in Hydra (Technau and Steele, 2011).

Here, we have established a novel approach which is based on previous RNAi applications in Hydra (Bosch et al., 2002; Lohmann and Bosch, 2000; Lohmann et al., 1999), but uses siRNAs for RNA interference (Watanabe et al., 2014). By refining the conditions for electroporation, we provide a robust protocol to silence target genes in Hydra with minimal tissue damage. Using our electroporation methodology, we also developed a reliable protocol for a CRISPR-Cas9 based knockout in adult animals. In our “somatic” CRISPR-Cas9 approach we directly deliver Cas9 ribonucleoproteins (RNPs) that are formed by the Cas9 protein and single-chain guide RNAs (sgRNAs) into adult polyps, thereby successfully introducing loss-of-function mutations in all three stem cell linages.

## Material and Methods

### Hydra Culture

Hydras were kept in Hydra medium (1 mM Tris pH 6.9, 1 mM NaHCO_3_; 0.1 mM KCl; 0.1 mM MgCl_2_; 1 mM CaCl_2_) at 18° C. Animals were fed three times a week with *Artemia salina* nauplii and Hydra medium was exchanged daily. The following Hydra strains were used: *Hydra vulgaris* AEP Act::GFP^ectoderm^/Act::RFP^endoderm^; *Hydra vulgaris* AEP Act::RFP^ectoderm^/Act::GFP^endoderm^ (Glauber et al., 2015), *Hydra vulgaris* AEP Hym-176::GFP nerve cells (Takahashi et al., 2008); *Hydra vulgaris* AEP CnNos1::GFP interstitial cells (Nishimiya-Fujisawa and Kobayashi, 2012).

### Electroporation of adult Hydra polyps

Medium-sized polyps were used that were fed one day before electroporation. Animals were collected with flame-polished glass pipettes in a Petri dish (Greiner BioOne) and shortly washed twice in ddH_2_O. 20 animals were then transferred into an electroporation cuvette (Gene Pulser / Micro Pulser Electroporation Cuvette 0.4 cm gap; Bio-Rad) and residual water was removed as completely as possible. siRNA was diluted in ddH_2_O to a final concentration of 1-5 μM and 200 μL of the solution was added to each cuvette. In case of Cas9-sgRNA ribonucleoproteins, a final concentration of 0.6 μM in 200 μl RNase free water was used, unless otherwise indicated. After relaxation of the animals, a single square wave pulse (200-300 V range) was applied for 10-50 ms (Gene Pulser XCell with CE Module; Bio-Rad). After electroporation, 500 μL of ice-cold recovery medium (80% hydra medium, 20% v/v dissociation medium: 3.6 mM KCl, 6 mM CaCl_2_, 1.2 mM MgSO4, 6 mM Na-Citrate, 6 mM pyruvate, 4 mM glucose, 12.5 mM TES pH6.9, 0.05 g/L rifampicin, 0.10 g/L streptomycin, 0.05 g/L kanamycin) was added to the animals, which were kept on ice until the experiment was completed. Animals were carefully transferred into a new Petri dish filled with pre-chilled recovery medium. It is important to note that excessive movement of the Petri dishes should be avoided during and after transfer. The polyps were left to recover overnight at 18 °C. On the next day, the recovery medium was exchanged for Hydra medium in a stepwise manner and debris was removed. Animals were fed daily from the second day of recovery. Visible gene silencing effects were detectable 3-4 days post-electroporation, peaked at day 6 and were stable until day 14, offering a range of 1 week for experiments.

### Small interfering RNA design

For the design of siRNAs, we followed general guidelines described previously (Birmingham et al., 2007;Shabalina et al., 2006). In general, we designed a 21 bp sequence starting with an adenine-dinucleotide, which is preferentially located in the 5’-region of the gene. Potential candidates were considered favorable when they contained a GC content between 30-50 % and were devoid of polyT sequences. Sequences were additionally refined by the prediction tool of the Whitehead Institute (http://jura.wi.mit.edu/bioc/siRNAext/). This tool offers the advantage of specificity prediction by considering thermodynamics. For each gene, three to four siRNAs were custom synthesized with an UU overhang (Sigma Aldrich) and tested for *in vivo* efficiency. siRNA specific for GFP was selected as follows: 5′-AACUACCUGUUCCAUGGCCAAUU-3′. A scrambled siGFP sequence was designed as control: 5′-AACUCAUCGAUUCACACCGGUUU-3′.

### RNA extraction and quantitative reverse transcription polymerase chain reaction

Ten animals that were electroporated with either siGFP or scrambled siGFP were randomly chosen and transferred into fresh 1.5mL tubes between day 7 and day 9 after electroporation. Residual Hydra medium was removed and animals were mixed with 500 μL TRIzol (Life Technologies) and homogenized by vortexing. After 5 minutes of incubation at RT, 100 μL chloroform (Sigma) was added to the suspension, thoroughly mixed and incubated for 5 minutes at RT followed by centrifugation at 12,000 xg for 15 minutes at 4 °C. The upper phase was transferred into a fresh 1.5 mL tube and mixed vigorously after adding 250 μL of chloroform:isoamyl-alcohol (24:1 v/v mixture; Sigma). After centrifugation at 12,000 xg for 15 minutes at 4 °C, the upper phase was transferred into a fresh 1.5 mL tube. In order to precipitate the RNA, the supernatant was mixed with 0.5 volumes of chilled isopropanol and incubated for 2 hours at −20 ^o^C followed by centrifugation as described above. The pellet was washed once with 70% ethanol (v/v), centrifuged at 12,000 x g for 15 minutes at 4 ^o^C and air-dried. The RNA was re-suspended in 30 μL ddH_2_O and subsequently digested with 1.5 U DNaseI (Roche) for 30 min at 37 °C. DNaseI was inactivated by heating the samples at 75 °C for 10 min. The concentration of the RNA was measured using a NanoDrop spectrophotometer. 1μg RNA per sample was employed for cDNA synthesis according to the manufacturer′s instructions (sensiFAST cDNA Synthesis Kit, Bioline). The cDNA was diluted 1:20 with 30μL ddH_2_O and used for quantitative PCR using SensiMix™ SYBR^®^ Hi-ROX Kit (Bioline), adopting the pipetting scheme and PCR conditions specified by the manufacturer. Primer pairs (MWG Eurofins) used were as follows: GFP (fw: 5′-TGATGCAACATACGGAAAACTTACC-3′; rv: 5′-ACACCAT AACAGAAAGTAGT GACAA-3′), elongation factor 1α as housekeeping gene for normalization (fw: 5′-TATTGATAGACCTTTTCGACTTTGC-3′; rv: 5′-CTGTAC AGAGCCACTTTCAACTTTT-3′). Samples were run in triplicates. Obtained data were analyzed using the ΔΔCt-method and visualized by Prism 7.0b (GraphPad).

### Purification of Cas9 protein

Purification of the Cas9 endonuclease was performed as described elsewhere (Gagnon et al., 2014). In brief, Cas9 protein was expressed in *E. coli* BL21(DE3) from the pET-28b-Cas9-His plasmid (Gagnon et al., 2014) via an auto-induction method (Studier, 2005). One liter of bacterial suspension was grown for 12 hrs at 37 °C, followed by a 24-hour expression at 20 °C. Cells were harvested, resuspended in washing buffer (20 mM Tris-HCl pH 8.0, 25 mM Imidazole, 500 mM NaCl), and lysed by sonication (duty cycle 20%, output control: 2,5; Branson Sonifier 250). The lysate was cleared by centrifugation for 15 minutes at 20,000 x g at 4 °C and loaded onto a 1 ml HisTrapHP column (GE-Healthcare). The column was washed with five column volumes of washing buffer and eluted using buffer containing 20 mM Tris-HCl pH 8.0, 500 mM Imidazole, 500 mM NaCl. 500 μl fractions were collected and the protein content was determined at 280 nm. Fractions with high protein concentration where pooled and dialyzed against a buffer containing 20 mM Tris-HCl pH 7.4, 200 mM KCl, 10 mM MgCl_2_. The final protein concentration was determined using the Pierce BCA Protein Assay Kit (Thermo Scientific) according to the manufacturer′s instructions. Single use aliquots were frozen in liquid nitrogen and stored at −80 °C.

### Single guide RNA design, sgRNA production and Cas9 *in vitro* activity assays

For the prediction of putative sgRNA binding sites the stand-alone version of the CCTop software (Stemmer et al., 2015) was used. The GFP coding sequence from pHyVec7 (GeneBank accession number: EF539830.1) served as an input sequence and the Hydra 2.0 genome assembly (https://arusha.nhgri.nih.gov/hydra/) was used as a reference genome. Among the predicted sgRNAs, the three top hits with the lowest off-target rate were chosen for further analyses. Single guide RNAs produced with the GeneArt Precision gRNA Synthesis Kit (Thermo Fisher Scientific) were as follows:

GFPsg1f (5′-TAATACGACTCACTATAGGGTGAAGGTGATGCAACATA-3′),

GFPsg1r (5′-TTCTAGCTCTAAACTATGTTGCATCACCTTCACC-3′),

GFPsg2f (5′-TAATACGACTCACTATAGTCAAGAGTGCCATGCCCGA-3′),

GFPsg2r (5′-TTCTAGCTCTAAACTCGGGCATGGCACTCTTGA-3′),

GFPsg3f (5′-TAATACGACTCACTATAGCATGCCGTTTCATATGATC-3′), and

GFPsg3r (5′-TTCTAGCTCTAAAC GATCATATGAAACGGCATG-3′).

To determine the *in vitro* activity of these sgRNAs a portion of the GFP coding sequence containing the sgRNA binding sites was amplified by PCR using primers GFPfwd1 (5′-GGAGAAGAACTTTTCACTGGA-3′) and GFPrev (5′-GTGTCCAAGAATGTTTCCAT-3′) to obtain a suitable cleavage template. *In vitro* cleavage reactions contained 100 mM NaCl, 5 mM MgCl_2_, 20 mM HEPES pH 6.5, 0.1 mM EDTA, 30 nM Cas9 nuclease, 30 nM sgRNA and 3 nM of DNA substrate. Reactions were incubated for 20 minutes at 37 °C. Cleavage reactions were then analyzed by agarose gel electrophoresis.

### Western Blot analysis

For protein extraction, polyps were resuspended in 2x sample buffer (Laemmli, 1970) at 10 μl per polyp and incubated at 95°C for 10 min with agitation. Protein extracts were cleared by centrifugation at 14,000 x g for 10 min, 0.1 hydra equivalents were loaded on 12 % SDS polyacrylamide gels, and after electrophoresis transferred to nitrocellulose membranes. Immunoblots were incubated with polyclonal chicken anti-GFP (1:5000, ab 13970; Abcam) and monoclonal mouse anti-a-tubulin (1:1000, clone DM1A, Sigma) primary antibodies and horseradish-peroxidase conjugated secondary rabbit anti-chicken (1:5000; 31401; Thermo Scientific) and goat anti-mouse (1:5000; 111689; Jackson ImmunoResearch,) antibodies. The blots were developed using the ECL system and bands were visualized on an ECL Chemocam Imager (Intas). Densitometric analysis of protein bands was performed using the Fiji software (Schindelin et al., 2012).

### Extraction of genomic DNA and mutation analysis

To extract genomic DNA, ten animals were placed in buffer containing 400 mM Tris-HCl pH 8.0, 5 mM EDTA pH 8.0, 150 mM NaCl, 0.1% Tween20, and 1 mg/ml Proteinase K and incubated at 60 °C overnight. The samples were extracted once with TE-saturated phenol, twice with diethyl ether and the genomic DNA was precipitated with ethanol following standard procedures. The GFP locus was amplified from genomic DNA by PCR using primers GFPfwd1 and GFPrev. The resulting fragment was subjected to a T7-Endonuclease analysis using the EnGenTM Mutation Detection Kit (New England Biolabs) following the manufacturer′s instructions. For sequence track decomposition analysis, a fragment of the GFP locus was amplified by PCR using primers GFPfwd2 (5′-AGTGGTTCACTGTACGTAA-3′) and GFPrev and used for Sanger sequencing (MWG Eurofins).The resulting sequence tracks were used for decomposition analysis using the TIDE software (Brinkman et al., 2014) (https://tide-calculator.nki.nl).

### Imaging

Pictures were taken between day 7-9 after electroporation, if not otherwise indicated. Animals were sedated in Chloretone (0.1%w/v in hydra medium) for live imaging. Fluorescent images of polyps were taken with a Nikon SMZ25 microscope equipped with a DS-RI2 camera and processed using NIS-Elements, Fiji, and Photoshop.

## Results and Discussion

The objective of this study was to present a robust electroporation protocol for reverse genetic methods in Hydra. In a first approach, we used transgenic Hydra strains expressing GFP under the control of actin or a cell type specific promotor and evaluated parameters such as animal viability and transfection efficiency employing siRNA specific to the introduced *GFP* gene (siGFP) and a corresponding scrambled sequence as control (scr-). We used transgenic strains that specifically express the GFP in the ectoderm or endoderm (Glauber et al., 2015), the interstitial stem cell lineage (Nishimiya-Fujisawa and Kobayashi, 2012), and the nervous system (Takahashi et al., 2008). Based on these results, we established in a second approach a protocol for a CRISPR-Cas9 based knockout by delivering Cas9 ribonucleoproteins (RNPs) into adult polyps. The induction of loss-of-function mutations was evaluated in both germ layers as well as in the interstitial stem cell lineage and differentiated neurons.

### Gene silencing by electroporation of siRNA

#### Effect of voltage and siRNA concentration on the viability of intact polyps

Previous publications showed a square wave pulse to be most efficient for introducing small molecules into cells with low transfection rates, such as JURKAT cells (Jordan et al., 2008). Accordingly, the square wave pulse was preferred to the exponential decay pulse in our initial experiments. Using this pulse pattern, animals were electroporated at increasing voltages as well as siRNA concentrations and were then evaluated for their general morphology and viability. Immediately after electroporation, the tissue of the animals exhibited a visibly reduced integrity in both cell layers, followed by a complete or partial loss of the tentacles (Fig. 1A). The transfer of the pulsed animals from the cuvettes to Petri dishes is a critical step and should be carried out carefully to avoid additional mechanical stress. In general, the integrity of the animals was restored 24 hours post electroporation. While animals pulsed at 200 V and 250 V or with a concentration of 1 μM and 3 μM siRNA showed moderate damage, polyps treated with either 300 V or 5 μM siGFP exhibited more severe injuries. Accordingly, Hydra pulsed at low or medium voltages and siRNA concentrations completely recovered within two days post electroporation with viability rates ranging from 94-100% (Fig. 1C). In contrast, polyps electroporated at 300 V or with 5 μM siGFP showed a more severe damage resulting in a decrease of viability to 70% and 84,5%, respectively. From these experiments, we conclude that 300 V and 5 μM siGFP represent the upper limits as electroporation parameters. We next examined whether these conditions were sufficient to induce an efficient silencing of the GFP mRNA.

**Fig. 1.**
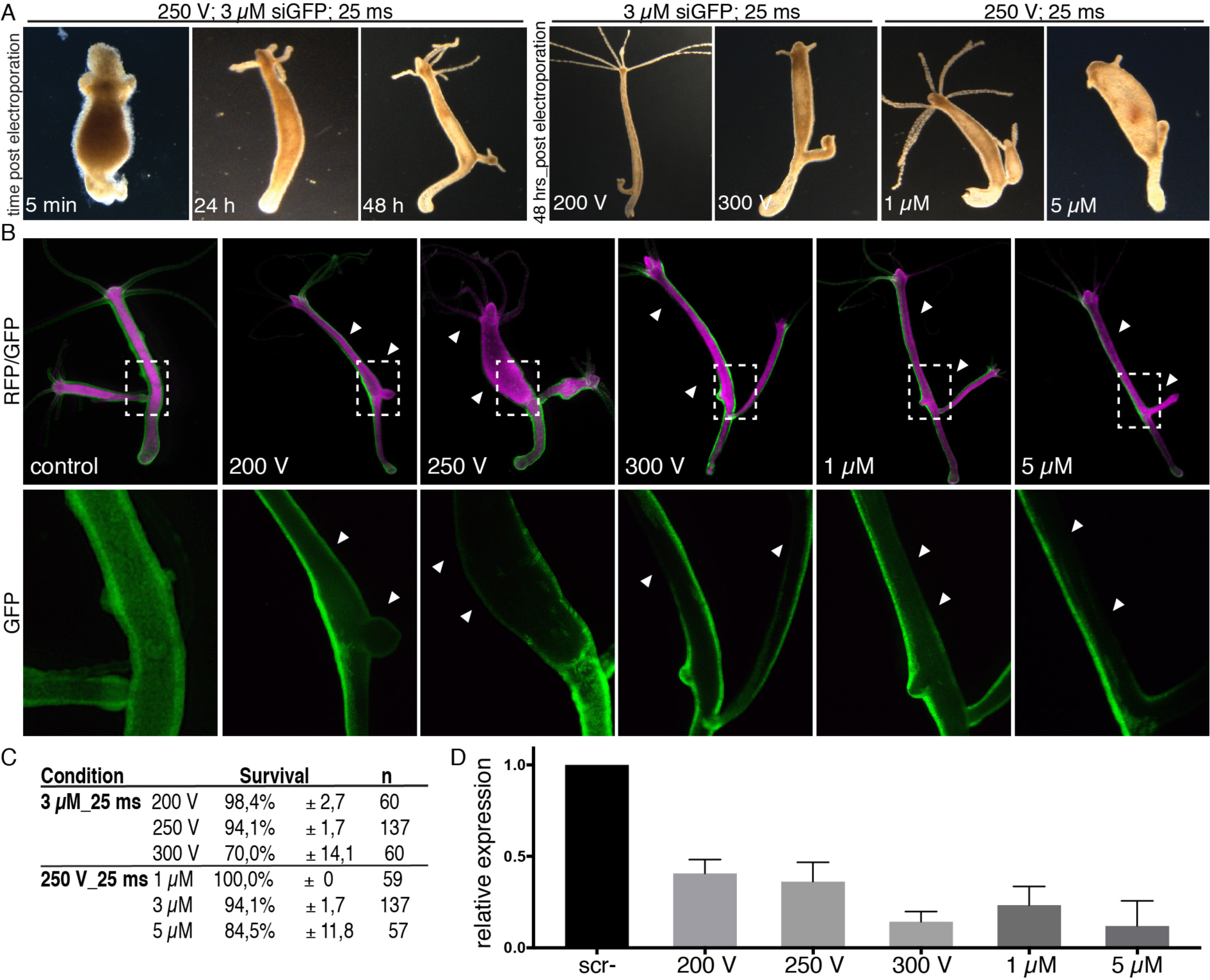
RNAi efficiency in dependence of voltage and siRNA concentration. (A) Animals pulsed at 200 V to 250 V or at a concentration of 1 μM siGFP (or less) recover within 48 hrs; polyps electroporated at 300 V or at a concentration of 5 μM siGFP do not show complete recovery. (B) Animals electroporated with siGFP exhibited a reduced fluorescence at the side of electroporation; scrambled control siRNA did not show any effects on GFP fluorescence. The silencing efficiency increases with higher voltages or application of higher siRNA concentrations. Upper panel shows whole polyps with both channels, GFP (green) and dsRed (magenta). Dashed rectangles mark the region used for magnification displayed in the lower panel. (C). The viability ranges between 94-100% for moderate electrotransfer conditions, while the application of high siRNA concentrations and high voltages results in survival decrease. (D) Quantitative analysis by RT-qPCR of whole animals confirms the reduction of GFP expression level. The degree of decrease is dependent on the stringency of the chosen electroporation conditions.

#### Effect of voltage on the knockdown efficiency

Electroporation at 200 V for 25 ms using 3 μM siGFP already showed a visible reduction of the GFP fluorescence starting from day 4 in the ecto::GFP animals, while the morphology remained completely unaffected (Fig. 1 A+B). In fact, animals pulsed under these conditions completely recovered within the first day after electroporation. Interestingly, the silencing of GFP was detectable only on one side of the animal, while the opposite side still possessed GFP fluorescence, though with a lesser intensity compared to the control. At higher magnification it became evident that the signal deriving from GFP was strongly reduced on the affected side, but not completely absent. A further decline in GFP fluorescence was achieved at a voltage of 250 V using the same concentration and pulse length. The animals exhibited a pronounced silencing of GFP, while still possessing a viability rate of 94% (Fig.1 C). The knockdown effect was additionally confirmed by RT-qPCR, which showed a decrease of the GFP expression level to almost one third relative to the scrambled control (Fig. 1D). It is important to note that RT-qPCR was performed with extracts deriving from whole animals that exhibit silencing only on one side of the animal. Thus, the actual decrease in the affected tissue area might be much higher. The use of 300 V reduced the expression level to 14% of the original amount illustrated by silenced areas of the animal lacking GFP fluorescence (Fig. 1B+D). However, this efficient reduction is associated with an increase of cell damage, morphological abnormalities, and a decrease of viability.

We next evaluated to which extent varying siRNA concentrations contributed to the knockdown efficiency. To this end, concentrations of 1 μM, 3 μM, and 5 μM siGFP were pulsed at constant 250 V for 25 ms. Although animals pulsed with 1 μM siGFP exhibited a reduction of the fluorescence comparable to polyps electroporated at 200 V, the expression of the GFP mRNA level dropped to 25% (Fig. 1B+D). This reduction on the transcriptional level is therefore in the range of an electroporation with 3 μM siGFP at 250 V for 25 ms. In contrast, the application of 5 μM induced a significant reduction of GFP mRNA to 12% as evaluated by RT-qPCR. However, animals pulsed under these conditions showed the most severe cell damage (Fig.1A-D).

All conditions tested resulted in silencing of the GFP expression at varying efficiencies. Taking together, a single square wave pulse at 250 V for 25 ms with a siGFP concentration of 3 μM allows for an efficient gene knockdown with minimal tissue damage.

#### Gene silencing in the interstitial stem cell lineage and endoderm

As the settings of 250 V for 25 ms in combination with 3 μM siGFP stably induced gene silencing in the ecto::GFP strain (Fig. 2A+B), we next asked whether these conditions were also suitable to target other cell types. Therefore, we applied the protocol to transgenic animals expressing GFP in the endoderm, interstitial cells, and nerve cells.

**Fig. 2.**
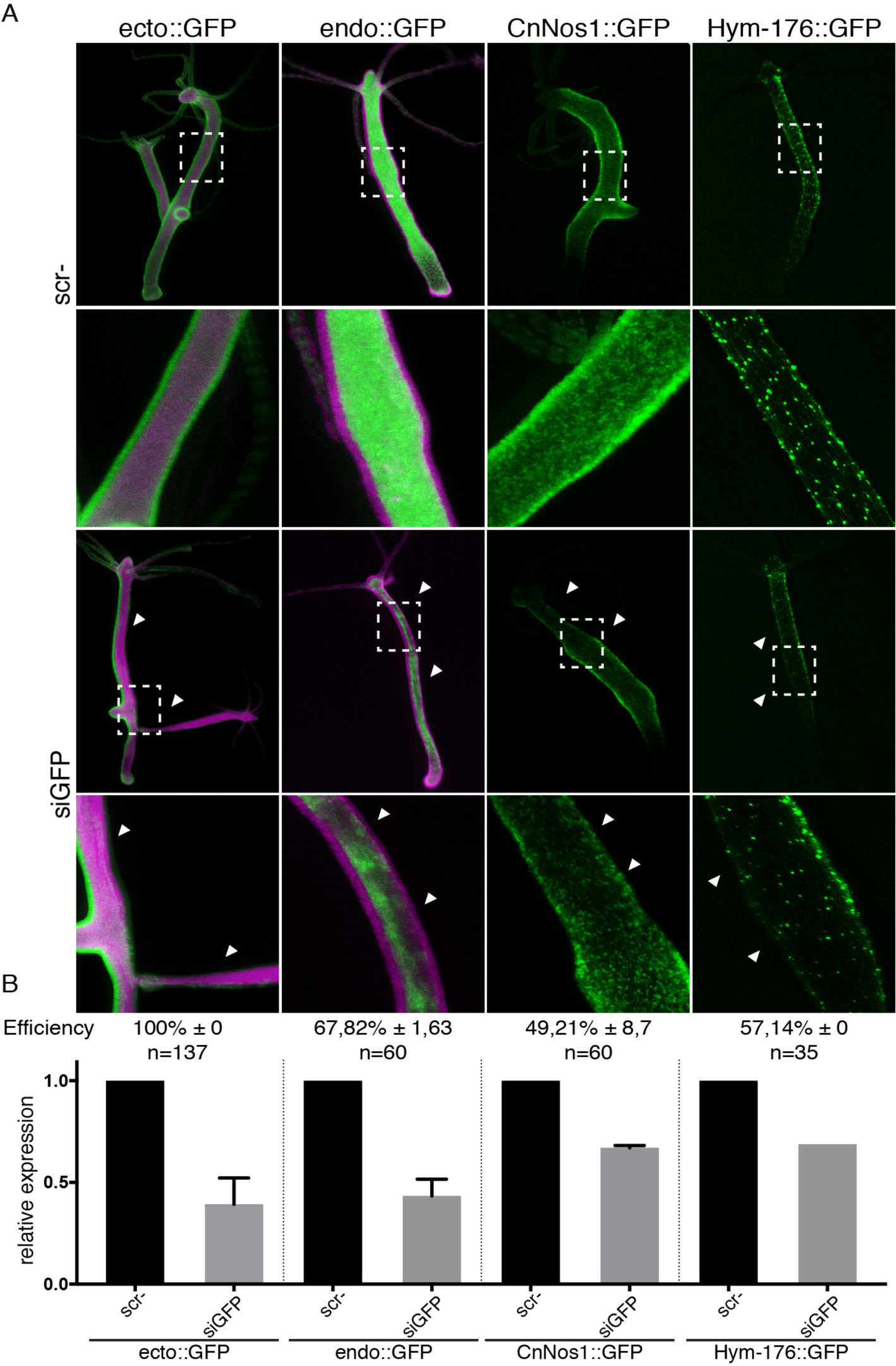
Targeting of different Hydra cell types. (A) Representative animals expressing GFP in the ectoderm (ecto::GFP, green), endoderm (endo::GFP, magenta), interstitial cells (CnNos1::GFP), and nerve cells (Hym-176::GFP). Dashed rectangles indicate areas displayed at higher magnification. A silencing of GFP can be visualized in all types with varying efficiencies (siGFP) Animals pulsed with scrambled siGFP (scr-) do not show a reduction in fluorescence. (B) The efficiency is expressed as percentage of animals exhibiting a reduction of GFP fluorescence relative to the total number of polyps pulsed. Quantitative analysis of whole animals by RT-qPCR reveals a reduction of the expression level ranging from 39-76% compared to the scrambled siGFP control.

Electroporation to the endo::GFP strain resulted in GFP silencing in 67% of the treated animals. Interestingly, the silenced tissue areas were distributed in a more scattered manner compared to the ecto::GFP strain that exhibited silencing predominantly on one side of the animal. However, the reduction of the total GFP mRNA levels with 43% was slightly lower compared to the ecto::GFP transgenic animals (Fig. 2B).

Since the interstitial cells of Hydra are rather small in size and moreover embedded within the ectodermal and endodermal layers, it has been very challenging to target these cells by electroporation (Smid and Tardent, 1984). To evaluate whether our electroporation settings were also applicable for the interstitial cell lineage, we performed GFP silencing in interstitial cell specific CnNos1::GFP transgenic animals. Using the settings already applied for the ecto::GFP and endo::GFP transgenic animals, we observed GFP silencing in 49% of the treated polyps. The moderate decrease of the transcript level to 67% was likely due to the fact that the affected tissue was smaller than in both epithelial germ layers. Comparable results were obtained with transgenic animals harboring GFP-labeled nerve cells (Hym-176::GFP). Given that 57% of the population exhibited a decrease in GFP expression, the efficiency is slightly higher compared to CnNOS1::GFP labeled animals, while the transcription level was reduced to a similar extent (Fig. 2A+B).

We can therefore conclude that the refined electroporation protocol enables an efficient silencing of GFP for two weeks (Fig. S1) in all cell types of Hydra.

### Genome editing by the electroporation of Cas9 guide RNA complexes in adult polyps

Although electroporation of siRNAs resulted in efficient reduction of transcript levels, duration of the knockdown was restricted to approximately 14 days, limiting phenotypic analyses. To overcome this issue, we aimed to develop an approach to directly interfere with gene function on a genomic level. With the emergence of programmable nucleases like transcription activator-like effector nucleases (TALEN (Miller et al., 2011)), Zinc-finger nucleases (ZNF (Urnov et al., 2010)), or RNA-guided nucleases, targeted genome editing has become available to virtually any model organism (Chen et al., 2013; Gratz et al., 2013; Hwang et al., 2013; Zhang et al., 2014). Among these, the clustered regularly interspaced short palindromic repeats (CRISPR)/CRISPR associated 9 (Cas9) system has become the method of choice. Here, the Cas9-nuclease is targeted to a specific genomic locus by a short single guide RNA (sgRNA) (Jinek et al., 2012). The Cas9-mediated introduction of a double strand break (DSB) at the genomic locus triggers non-homologous end joining, an error-prone cellular repair mechanism, that leads to the insertion and/or deletion (InDels) at the DSB (Gallagher and Haber, 2017).

### Basic protocol for the electroporation of Cas9 guide RNA complexes

Previous studies have demonstrated that ribonucleoproteins of Cas9 protein and sgRNA can be introduced into cells by electroporation, suggesting that this method might be suitable for genome editing in adult Hydra polyps. To test this hypothesis, we first aimed to target the recombinant GFP locus of ecto::GFP animals using three independent CRISPR target sites (Fig. 3A and Material and Methods). DNA cleavage by the corresponding guide RNAs was confirmed by an *in vitro* Cas9 cleavage assay using a PCR-fragment derived from the genomic GFP locus (Fig. 3B). We added Cas9 protein to the individual GFP-sgRNAs in a 1:1.2 molar ratio to allow the formation of ribonucleoproteins (Cas9RNP) and used these complexes for the electroporation experiment. Using a concentration of 0.6 μM Cas9RNP the animals were subjected to a single 25 ms pulse at 230 V. Effects on GFP expression were determined seven days after electroporation. While animals treated with Cas9 protein only, Cas9-GFPsg2 or Cas9-GFP RNPs did not show any obvious phenotype, we observed areas with absent GFP signal in polyps that were electroporated with Cas9-GFPsg1 RNPs (Fig. 3C). This loss of GFP signal correlated with sequence aberrations at the GFP-locus as detected by a T7 endonuclease assay. Mismatch cleavage by the T7 endonuclease was only detectable on DNA heteroduplexes derived from animals that were electroporated with Cas9-GFPsg1 RNPs (Fig. 3D). Quantitative assessment of genome editing by sequence trace decomposition using the TIDE software (Brinkman et al., 2014) indicated mosaic editing of the GFP coding sequence in these polyps. Overall, 29.1% of the DNA was edited, resulting in a four-base deletion in most cases (Fig. 3E). Finally, we assessed GFP protein levels in targeted animals. Quantitative immunoblot analyses revealed a slight reduction in GFP expression in 16% of Cas9-GFPsg1 RNP electroporated animals compared to the control. By comparison, protein levels in Cas9-GFPsg2 RNP and Cas9-GFPsg3 RNP treated animals remained unaffected (Fig. 3F). Taken together, these results demonstrate that CRISPR/Cas9 mediated genome editing in adult hydra polyps can be induced by the electroporation of Cas9-sgRNA RNPs.

**Fig. 3.**
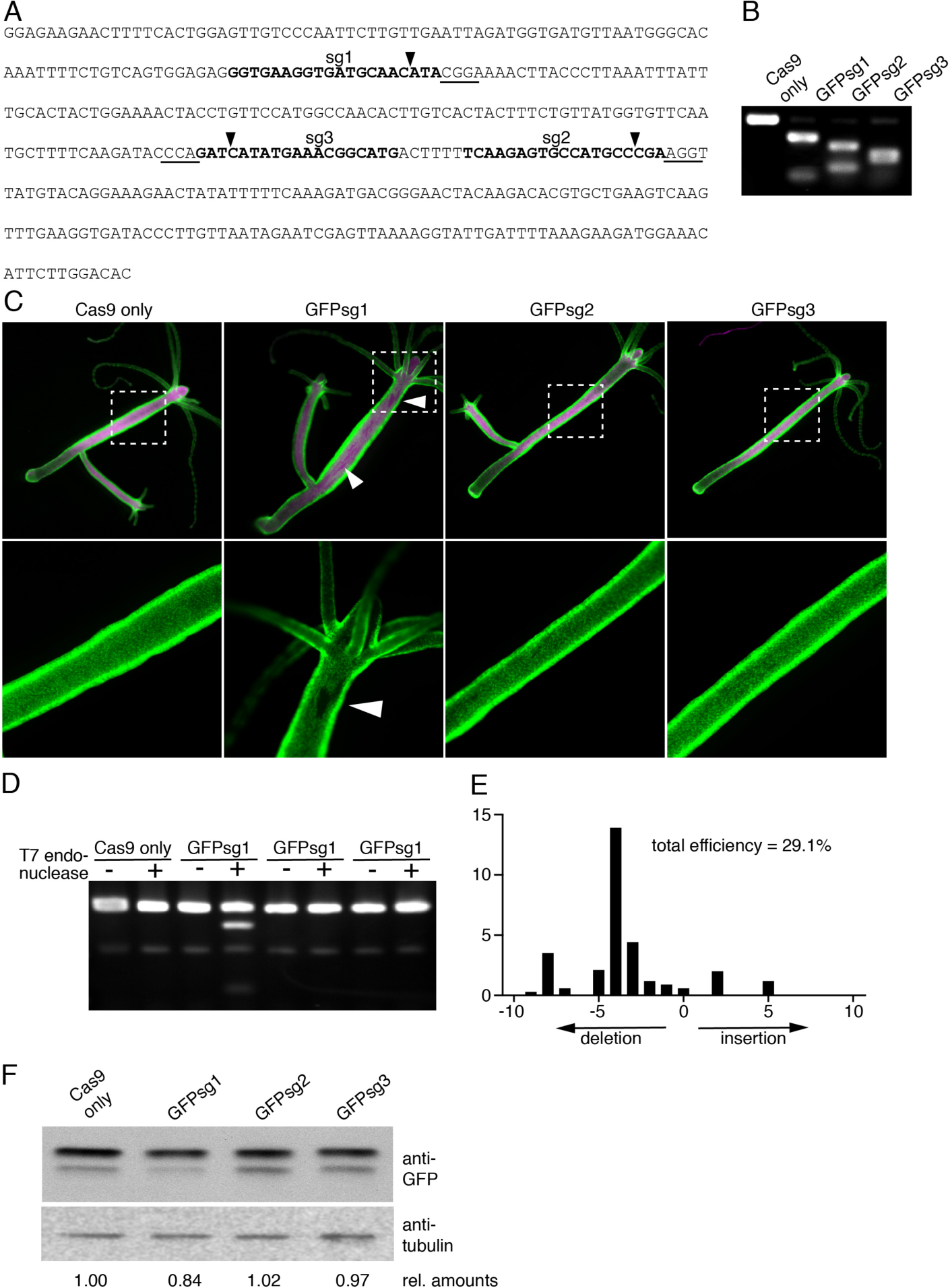
Electroporation of Cas9 guide RNA complexes results in genome editing in adult polyps. (A) Partial representation of the GFP coding sequence. Target sites of sgRNAs used are shown in bold. Corresponding protospacer adjacent motifs are underlined. Arrowheads indicate cleavage sites of Cas9-sgRNPs. (B) *In vitro* Cas9 cleavage assay. A PCR fragment corresponding to the sequence in (A) was incubated with either Cas9 only (control) or Cas9 in combination with the sgRNAs GFPsg1-3. Presence of individual sgRNAs led to the anticipated cleavage products (GFPsg1: 104 bp + 304 bp, GFPsg2: 150 bp + 257 bp, GFPsg3: 189 bp + 219 bp). (C) Fluorescence microscopy of polyps treated with either Cas9 protein or Cas9 in combination with corresponding sgRNAs, as indicated, 7 days after electroporation. Upper panel shows an overview of the whole polyp. Superimpositions of GFP (green) and dsRed (magenta) channels are shown. A dashed rectangle indicates the area shown at high magnification in the lower panel. Arrowheads in Cas9-GFPsg1 RNP electroporated animals indicate regions with absent GFP signal. (D) Mismatch cleavage assay in Cas9 only and Cas9-GFPsg1-3 treated animals. PCR fragments of the GFP locus (A) were denatured and reannealed to allow heteroduplex formation (wildtype and mutated DNA strand). Heteroduplexes were interrogated by T7 endonuclease that cuts imperfectly matched DNA. Arrowheads indicate cleavage products in Cas9-GFPsg1 electroporated animals at 100 and 300bp. (E) Quantitative assessment of InDels in the GFP coding sequence of Cas9-GFPsg1 treated polyps. The GFP locus of treated and mock-treated animals was sequenced by Sanger sequencing following decomposition of the sequence tracks using the TIDE software (Brinkman et al., 2014). The relative abundance of insertion and deletions of every length with respect to the Cas9-sgRNA RNP cutting site are shown. (F) Protein extracts from treated and mock-treated animals were analyzed using GFP-specific antibodies to assess GFP protein levels. Tubulin served as a loading control. Numbers indicate the relative amounts of GFP protein that were determined by densitometric analysis of the bands. Tubulin signals were used to normalize loading.

### Optimized conditions for RNP electroporation

Although we clearly detected gene targeting in our electroporated animals, genome editing efficiencies were relatively low. Thus, we next aimed to optimize our protocol for the delivery of the Cas9-sgRNA RNPs. To that end, we tested different concentrations of the Cas9-sgRNA RNP as well as various voltages in our electroporation assay and assessed the gene targeting efficiency by analyzing genome editing of the GFP locus, GFP protein levels, and survival rates. Conditions were varied by using either 0.3 μM to 1.2 μM of Cas9-RNPs at a constant voltage of 230 V or 210 V to 270 V at a constant concentration of Cas9-RNPs. A maximum of GFP depleted areas was observed in polyps that were treated with 0.6 μM Cas9-GFPsg1 RNP or pulsed at 250 V, respectively (Fig. S2 A and D). This observation was confirmed by the quantitative data we obtained. In the presence of 0.6 μM RNPs or 250 V we obtained a genome editing efficiency of approximately 70% in nearly 80% of the animals, which resulted in a 50% decrease in GFP protein levels (Fig. S2 B, C, E, and F). Higher RNP concentration or a further increase in the voltage applied had only little or no effect on the targeting efficiencies but was detrimental to the survival rates after electroporation (Fig S2 B and E). In summary, these results suggest optimal electroporation conditions at 250 V in the presence of 0.6 μM of the Cas9 sgRNA complex. These conditions resulted in high genome editing rates while sustaining sufficient amounts of surviving animals for downstream analysis. Moreover, after the initial recovery phase, the treated polyps showed little or no morphological abnormalities that would obscure phenotypic read-outs.

### Electroporation of Cas9-sgRNA RNP mediates genome editing in all stem cell lineages

So far, we had restricted our analysis to the ectoderm of the hydra polyp. In order to further extend this study to additional cell types we performed electroporation using the aforementioned transgenic animals that recombinantly express GFP either in the endoderm, the interstitial cells, or nerve cells. Fluorescence microscopy analysis of the treated polyps revealed areas devoid of GFP signal in the majority of all animals tested (Fig. 4A). Genome editing was further confirmed by a mismatch cleavage assay using the T7 endonuclease and decomposition analysis revealing that 50-70% of the GFP loci were targeted (Fig. 4B and C). Moreover, we observed a clear decrease of GFP protein levels in the treated animals compared to mock-treated controls (Fig. 4D).

**Fig. 4.**
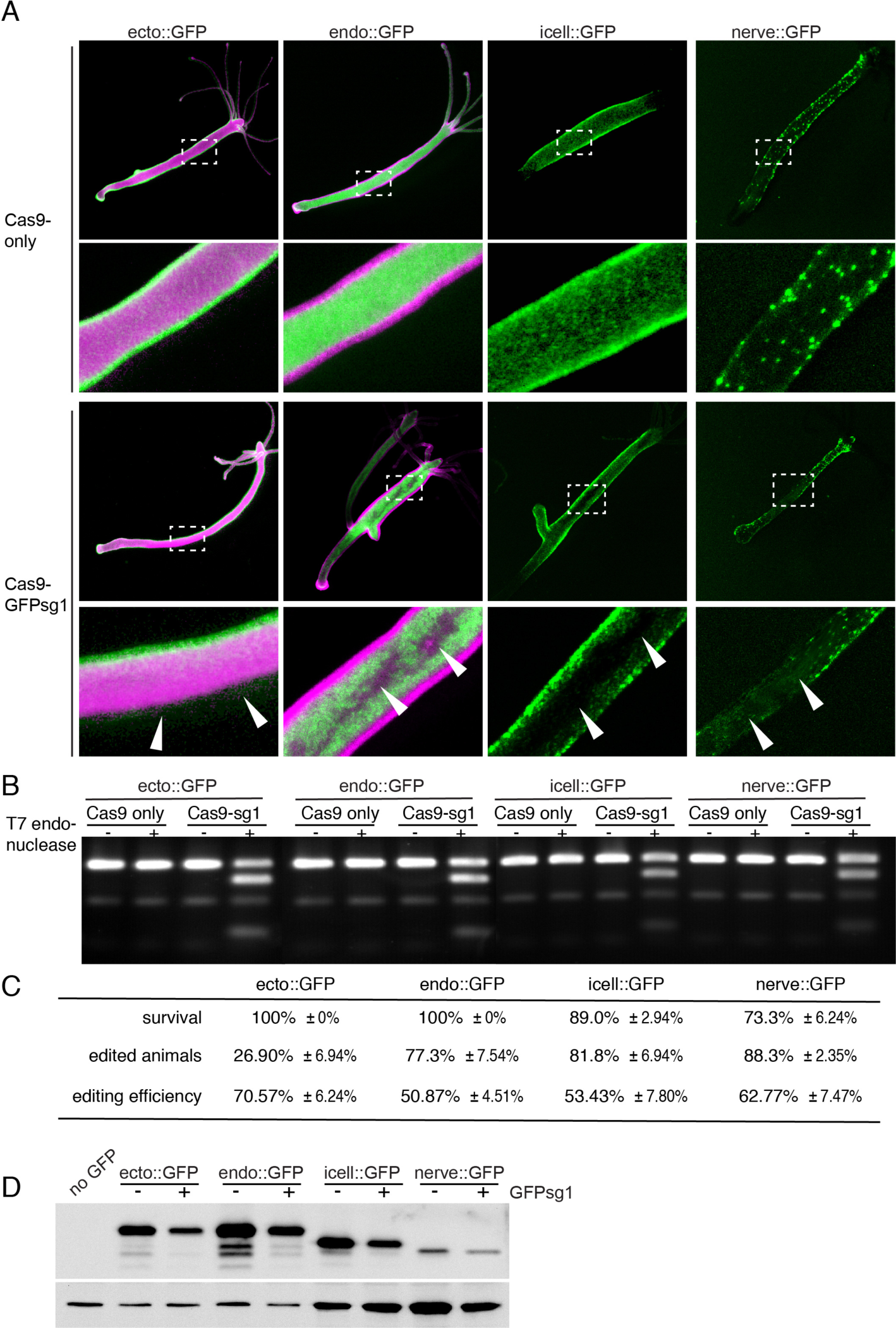
Electroporation of Cas9-sgRNA RNP mediates genome editing in all stem cell lineages. (A) Fluorescence microscopy of hydra strains featuring GFP expression in the ectoderm, the endoderm, the interstitial cells, and in nerve cells as indicated. Animals were electroporated with either Cas9 proteins only or with Cas9-GFPsg1 RNPs and analyzed 7 days after electroporation. Superimpositions of the GFP-(green) and the dsRed (magenta) channel are shown. The dashed rectangle in the upper panel indicates the area that is shown at high magnification in the lower panel. Arrowheads indicate regions devoid of GFP signal in the Cas9-GFPsg1 RNP-treated animals. (B) Mismatch cleavage assay of treated and mock-treated polyps from the different Hydra lines analyzed. See caption of Fig. 3 for further details. (C) Quantitative assessment of surviving animals, amounts of animals showing altered GFP expression, and genome editing efficiency obtained by TIDE analysis. Average numbers from three independent experiments are shown. (D) GFP protein levels in the electroporated animals. Western Blot analyses using anti-GFP antibodies were performed to visualize differences in GFP protein amounts. After analysis with the anti-GFP antibodies, tubulin was visualized as a loading control.

Taken together, these results demonstrate that CRISPR/Cas9-mediated genome editing after electroporation of Cas9-sgRNA RNPs into adult Hydra polyps is not restricted to ectodermal tissue but also allows targeting of endodermal and interstitial cells. Since the mutations generated by CRISPR/Cas9-mediated genome editing are stably integrated into the genome it should be possible to generate stable mutant lines for all three stem cell populations in Hydra. To test this hypothesis, we excised tissue parts devoid of GFP signal from an ecto::GFP animal electroporated with Cas9-GFPsg1 RNPs (supplemental Fig. S3). The resulting polyps from these regenerates showed a minimal number of GFP positive ectodermal cells and produced buds completely devoid of GFP expression (data not shown). Up to date we have cultured these polyps for more than a year without noticing any reappearance of GFP expression (supplemental Fig. S3).

Our approach demonstrates that CRISPR/Cas9-mediated genome editing can be used to introduce stable mutations in the genome of Hydra polyps. For functional knockout studies it will be necessary to target both alleles of an endogenous gene. Here, we have not investigated if such bi-allelic events are induced since our study is restricted to a transgene that was introduced into the genome of polyps by microinjection of plasmid DNA into Hydra embryos. However, a recent study has shown that such transgenes often integrate into genomes at multiple sites and with multiple copies (Wittlieb et al., 2006). Thus, it is possible that the transgenic animals used in this study also bear multiple copies of the GFP-transgene. In that case, the regions devoid of GFP expression in our treated animals might contain cells, in which multiple loci were targeted. Further experiments will be needed to confirm this assumption.

### Perspective

For many years of research in Hydra developmental biology was dominated by the analysis of gene expression patterns. These studies were extremely useful in collecting a large database for comparative analyses of major signaling pathways and transcriptions factors between cnidaria and bilateria. However, mechanistic studies were largely restricted to the application of pharmacological treatments with often questionable specificity. We expect that our robust protocols for siRNA knockdowns and CRISPR/Cas9-based genome editing will open new avenues for functional gene analysis in Hydra and yield important insights into basic mechanisms of stem cell and regeneration biology.

## Acknowledgements

We thank R. Steele and C. Fujisawa for providing transgenic Hydra strains. This study was supported by the Heidelberg Excellence Cluster Cellular Networks and grants from the German science foundation (DFG) to T.W.H. (SFB 873/A1; SFB 1324/A5). We thank Suat Özbek for many fruitful discussions and critically reading the manuscript.

## Author Contributions

A.T., M.L. and T.W.H. designed the research. A.T., M.L. B.T. and L.R.-C. performed experiments. A.T., M.L., and T.W.H. wrote the manuscript.

---

**Fig. S1.**
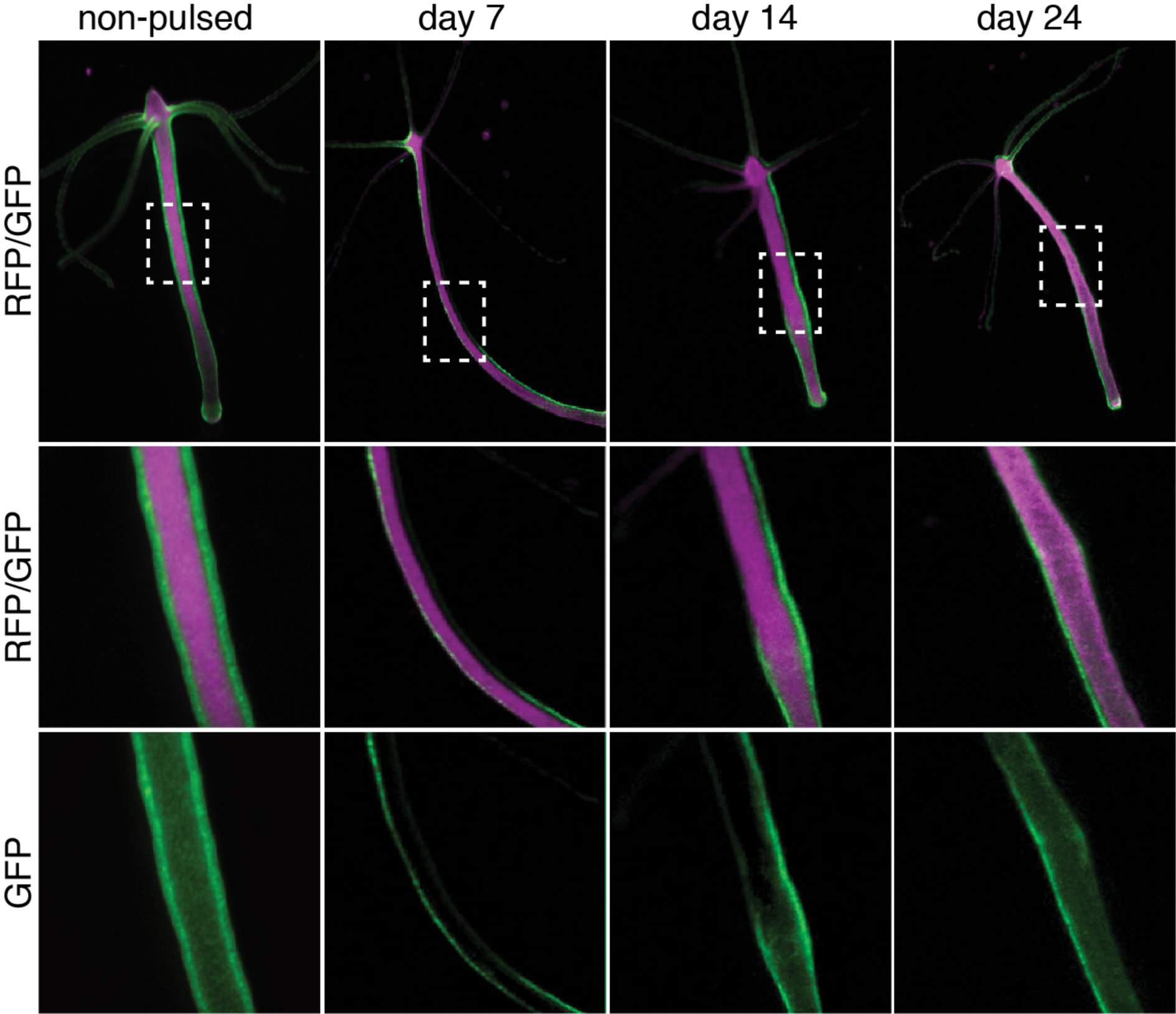
Stability of siRNA-mediated knockdown of GFP. The silencing of the GFP gene lasted for 14 days. GFP fluorescence started to return after approximately 3 weeks.

**Fig. S2.**
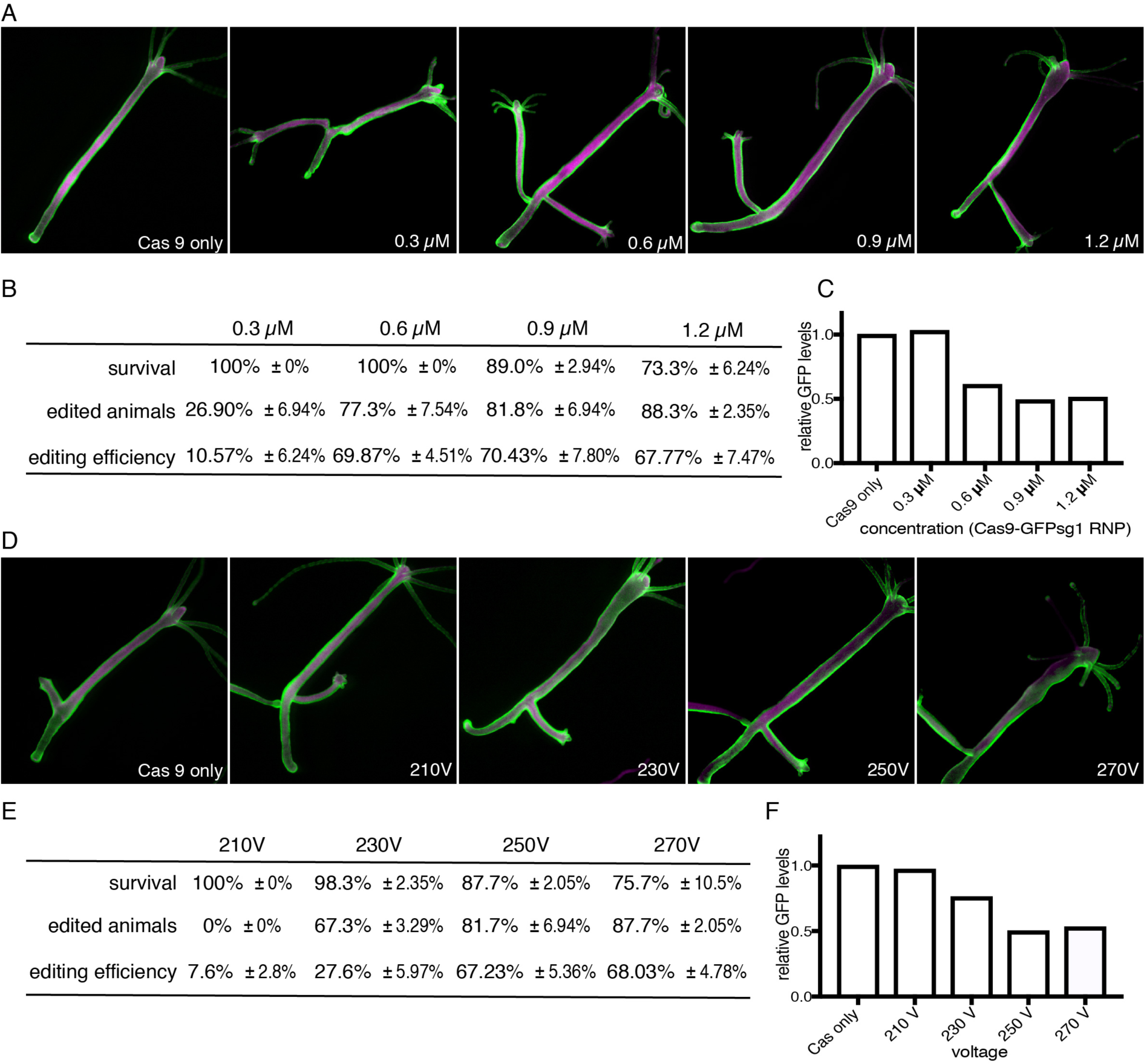
Optimization of Cas9-sgRNA delivery in adult polyps. (A and D) Fluorescence microscopy of Hydra polyps electroporated with different Cas9-GFPsg1 RNP concentrations or at different voltages as indicated. Superimposed images of the GFP (green) and the sdRed (magenta) channels are shown. (B and E) Quantitative assessment of the animal survival, number of edited animals, and genome editing efficiency (according to TIDE analysis) of polyps electroporated using the indicated conditions. Mean values from three independent experiments are shown. (C and F) Quantitative analysis of GFP protein levels in animals electroporated under different conditions compared to mock-treated animals. The relative amounts of GFP protein were determined by quantitative Western Blot analysis. Protein amounts in the different extracts were normalized using tubulin.

**Fig. S3.**
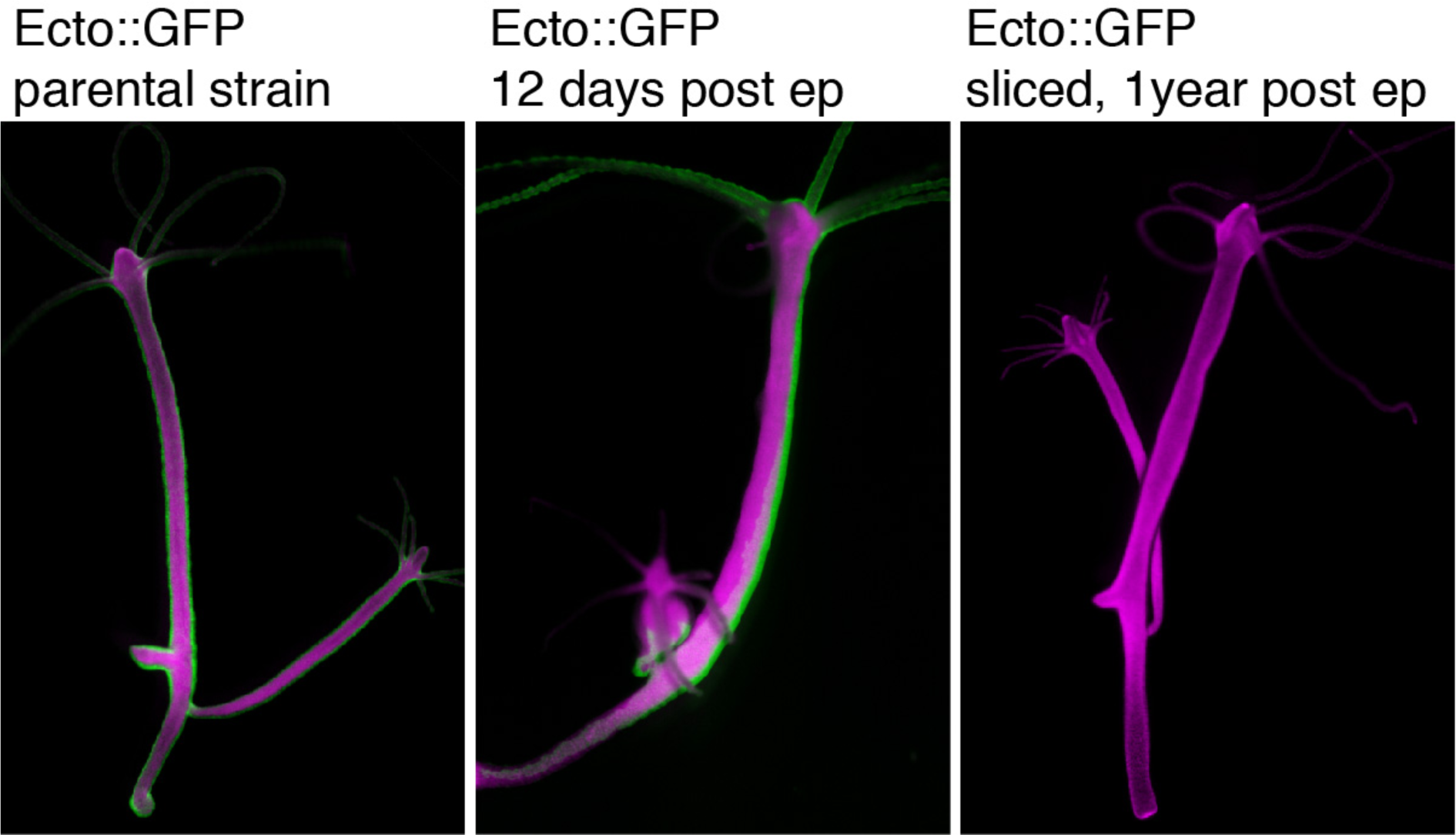
Generation of stable GFP-mutant lines by CRISPR/Cas9 mediated genome editing. Polyps of the ecto::GFP Hydra strain (left) were electroporated with Cas9-GFPsg1 RNPs to induce genome editing at the GFP-locus. Twelve days post electroporation (middle), tissue regions devoid of GFP expression were excised and allowed to regenerated into intact polyps, which produced offspring completely lacking GFP expression. This GFP-less phenotype was stably maintained even after one year of culture (right).

## References

Birmingham, A., Anderson, E., Sullivan, K., Reynolds, A., Boese, Q., Leake, D., Karpilow, J., Khvorova, A., 2007. A protocol for designing siRNAs with high functionality and specificity. Nat Protoc 2, 2068–2078.

Bosch, T.C., Anton-Erxleben, F., Hemmrich, G., Khalturin, K., 2010. The Hydra polyp: nothing but an active stem cell community. Dev Growth Differ 52, 15–25.

Bosch, T.C., Augustin, R., Gellner, K., Khalturin, K., Lohmann, J.U., 2002. In vivo electroporation for genetic manipulations of whole Hydra polyps. Differentiation 70, 140–147.

Brinkman, E.K., Chen, T., Amendola, M., van Steensel, B., 2014. Easy quantitative assessment of genome editing by sequence trace decomposition. Nucleic Acids Res 42, e168.

Chen, C., Fenk, L.A., de Bono, M., 2013. Efficient genome editing in Caenorhabditis elegans by CRISPR-targeted homologous recombination. Nucleic Acids Res 41, e193.

Conte, D., Jr., Mello, C.C., 2003. RNA interference in Caenorhabditis elegans. Curr Protoc Mol Biol Chapter 26, Unit 26 23.

David, C.N., 2012. Interstitial stem cells in Hydra: multipotency and decision-making. Int J Dev Biol 56, 489–497.

Elbashir, S.M., Harborth, J., Lendeckel, W., Yalcin, A., Weber, K., Tuschl, T., 2001. Duplexes of 21-nucleotide RNAs mediate RNA interference in cultured mammalian cells. Nature 411, 494–498.

Fire, A., Xu, S., Montgomery, M.K., Kostas, S.A., Driver, S.E., Mello, C.C., 1998. Potent and specific genetic interference by double-stranded RNA in Caenorhabditis elegans. Nature 391, 806–811.

Gagnon, J.A., Valen, E., Thyme, S.B., Huang, P., Akhmetova, L., Pauli, A., Montague, T.G., Zimmerman, S., Richter, C., Schier, A.F., 2014. Efficient mutagenesis by Cas9 protein-mediated oligonucleotide insertion and large-scale assessment of single-guide RNAs. PLoS One 9, e98186.

Gallagher, D.N., Haber, J.E., 2017. Repair of a Site-Specific DNA Cleavage: Old-School Lessons for Cas9-Mediated Gene Editing. ACS Chem Biol.

Glauber, K.M., Dana, C.E., Park, S.S., Colby, D.A., Noro, Y., Fujisawa, T., Chamberlin, A.R., Steele, R.E., 2013. A small molecule screen identifies a novel compound that induces a homeotic transformation in Hydra. Development 140, 4788–4796.

Gratz, S.J., Cummings, A.M., Nguyen, J.N., Hamm, D.C., Donohue, L.K., Harrison, M.M., Wildonger, J., O′Connor-Giles, K.M., 2013. Genome engineering of Drosophila with the CRISPR RNA-guided Cas9 nuclease. Genetics 194, 1029–1035.

Holstein, T.W., 2012. The evolution of the Wnt pathway. Cold Spring Harb Perspect Biol 4, a007922.

Holstein, T.W., Watanabe, H., Ozbek, S., 2011. Signaling pathways and axis formation in the lower metazoa. Curr Top Dev Biol 97, 137–177.

Hwang, W.Y., Fu, Y., Reyon, D., Maeder, M.L., Tsai, S.Q., Sander, J.D., Peterson, R.T., Yeh, J.R., Joung, J.K., 2013. Efficient genome editing in zebrafish using a CRISPR-Cas system. Nat Biotechnol 31, 227–229.

Jinek, M., Chylinski, K., Fonfara, I., Hauer, M., Doudna, J.A., Charpentier, E., 2012. A programmable dual-RNA-guided DNA endonuclease in adaptive bacterial immunity. Science 337, 816–821.

Jordan, E.T., Collins, M., Terefe, J., Ugozzoli, L., Rubio, T., 2008. Optimizing electroporation conditions in primary and other difficult-to-transfect cells. J Biomol Tech 19, 328–334.

Laemmli, U.K., 1970. Cleavage of structural proteins during the assembly of the head of bacteriophage T4. Nature 227, 680–685.

Lohmann, J.U., Bosch, T.C., 2000. The novel peptide HEADY specifies apical fate in a simple radially symmetric metazoan. Genes Dev 14, 2771–2777.

Lohmann, J.U., Endl, I., Bosch, T.C.G., 1999. Silencing of developmental genes in Hydra. Developmental Biology 214, 211–214.

Miljkovic-Licina, M., Chera, S., Ghila, L., Galliot, B., 2007. Head regeneration in wild-type hydra requires de novo neurogenesis. Development 134, 1191–1201.

Miller, J.C., Tan, S., Qiao, G., Barlow, K.A., Wang, J., Xia, D.F., Meng, X., Paschon, D.E., Leung, E., Hinkley, S.J., Dulay, G.P., Hua, K.L., Ankoudinova, I., Cost, G.J., Urnov, F.D., Zhang, H.S., Holmes, M.C., Zhang, L., Gregory, P.D., Rebar, E.J., 2011. A TALE nuclease architecture for efficient genome editing. Nat Biotechnol 29, 143–148.

Momose, T., Derelle, R., Houliston, E., 2008. A maternally localised Wnt ligand required for axial patterning in the cnidarian Clytia hemisphaerica. Development 135, 2105–2113.

Nakamura, Y., Tsiairis, C.D., Ozbek, S., Holstein, T.W., 2011. Autoregulatory and repressive inputs localize Hydra Wnt3 to the head organizer. Proc Natl Acad Sci U S A 108, 9137–9142.

Newmark, P.A., Reddien, P.W., Cebria, F., Sanchez Alvarado, A., 2003. Ingestion of bacterially expressed double-stranded RNA inhibits gene expression in planarians. Proc Natl Acad Sci U S A 100 Suppl 1, 11861–11865.

Nishimiya-Fujisawa, C., Kobayashi, S., 2012. Germline stem cells and sex determination in Hydra. Int J Developmental Biology 56, 499–508.

Rentzsch, F., Fritzenwanker, J.H., Scholz, C.B., Technau, U., 2008. FGF signalling controls formation of the apical sensory organ in the cnidarian Nematostella vectensis. Development 135, 1761–1769.

Schaible, R., Scheuerlein, A., Danko, M.J., Gampe, J., Martinez, D.E., Vaupel, J.W., 2015. Constant mortality and fertility over age in Hydra. Proc Natl Acad Sci U S A 112, 15701–15706.

Schindelin, J., Arganda-Carreras, I., Frise, E., Kaynig, V., Longair, M., Pietzsch, T., Preibisch, S., Rueden, C., Saalfeld, S., Schmid, B., Tinevez, J.Y., White, D.J., Hartenstein, V., Eliceiri, K., Tomancak, P., Cardona, A., 2012. Fiji: an open-source platform for biological-image analysis. Nat Methods 9, 676–682.

Shabalina, S.A., Spiridonov, A.N., Ogurtsov, A.Y., 2006. Computational models with thermodynamic and composition features improve siRNA design. BMC Bioinformatics 7, 65.

Simion, P., Philippe, H., Baurain, D., Jager, M., Richter, D.J., Di Franco, A., Roure, B., Satoh, N., Queinnec, E., Ereskovsky, A., Lapebie, P., Corre, E., Delsuc, F., King, N., Worheide, G., Manuel, M., 2017. A Large and Consistent Phylogenomic Dataset Supports Sponges as the Sister Group to All Other Animals. Curr Biol 27, 958–967.

Smid, I., Tardent, P., 1984. Migration of I-cells from ectoderm to endoderm in Hydra attenuata Pall (Cnidaria, Hydrozoa) and their subsequent differentiation. Developmental Biology 106, 469–477.

Smith, K.M., Gee, L., Bode, H.R., 2000. HyAlx, an aristaless-related gene, is involved in tentacle formation in hydra. Development 127, 4743–4752.

Steele, R.E., 2002. Developmental signaling in Hydra: what does it take to build a “simple” animal? Developmental Biology 248, 199–219.

Steele, R.E., 2012. The Hydra genome: insights, puzzles and opportunities for developmental biologists. Int J Dev Biol 56, 535–542.

Steele, R.E., David, C.N., Technau, U., 2011. A genomic view of 500 million years of cnidarian evolution. Trends Genet 27, 7–13.

Stemmer, M., Thumberger, T., Del Sol Keyer, M., Wittbrodt, J., Mateo, J.L., 2015. CCTop: An Intuitive, Flexible and Reliable CRISPR/Cas9 Target Prediction Tool. PLoS One 10, e0124633.

Studier, F.W., 2005. Protein production by auto-induction in high density shaking cultures. Protein Expr Purif 41, 207–234.

Takahashi, T., Hayakawa, E., Koizumi, O., Fujisawa, T., 2008. Neuropeptides and their functions in Hydra. Acta Biol Hung 59 Suppl, 227–235.

Technau, U., Steele, R.E., 2011. Evolutionary crossroads in developmental biology: Cnidaria. Development 138, 2143–2152

Trembley, A., 1744. Mémoires pour servir à l′histoire d′un genre de polypes d′eau douce à bras en forme de cornes. J.& H. Verbeek, Leide.

Urnov, F.D., Rebar, E.J., Holmes, M.C., Zhang, H.S., Gregory, P.D., 2010. Genome editing with engineered zinc finger nucleases. Nat Rev Genet 11, 636–646.

Watanabe, H., Schmidt, H.A., Kuhn, A., Hoger, S.K., Kocagoz, Y., Laumann-Lipp, N., Ozbek, S., Holstein, T.W., 2014. Nodal signalling determines biradial asymmetry in Hydra. Nature 515, 112–115.

Wittlieb, J., Khalturin, K., Lohmann, J.U., Anton-Erxleben, F., Bosch, T.C., 2006. Transgenic Hydra allow in vivo tracking of individual stem cells during morphogenesis. Proc Natl Acad Sci U S A 103, 6208–6211.

Zhang, H., Zhang, J., Wei, P., Zhang, B., Gou, F., Feng, Z., Mao, Y., Yang, L., Zhang, H., Xu, N., Zhu, J.K., 2014. The CRISPR/Cas9 system produces specific and homozygous targeted gene editing in rice in one generation. Plant Biotechnol J 12, 797–807.

